# Comprehensive summary of steroid metabolism in *Comamonas testosteroni* TA441; entire degradation process of basic four rings and removal of C12 hydroxyl group

**DOI:** 10.1101/2023.02.02.526766

**Authors:** Masae Horinouchi, Toshiaki Hayashi

**Affiliations:** Environmental Molecular Biology Laboratory, RIKEN, Saitama, 351-0198 Japan; Surface and Interface Science Laboratory, RIKEN, Saitama, 351-0198 Japan

**Keywords:** *Comamonas testosteroni*, bile acid, cholic acid, testosterone, cholesterol, steroid, β-oxidation, brain-gut-microbiome axis, brain-gut axis, *Mycobacterium tuberculosis*

## Abstract

*Comamonas testosteroni* is one of the representative aerobic steroid-degrading bacterium. We previously revealed the mechanism of steroidal A,B,C,D-ring degradation by *C. testosteroni* TA441. The corresponding genes are located in two clusters at both ends of a mega-cluster of steroid degradation genes. ORF7 and ORF6 are the only two genes in these clusters, whose function has not been determined. Here, we characterized ORF7 as encoding the dehydrase responsible for converting the C12β hydroxyl group to the C10(12) double bond on the C-ring (SteC), and ORF6 as encoding the hydrogenase responsible for converting the C10(12) double bond to a single bond (SteD). SteA and SteB, encoded just upstream of SteC and SteD, are in charge of oxidizing the C12α hydroxyl group to a ketone group, and of reducing the latter to the C12β hydroxyl group, respectively. Therefore, the C12α hydroxyl group in steroids is removed with SteABCD via the C12 ketone and C12β hydroxyl groups. Given the functional characterization of ORF6 and ORF7, we disclose the entire pathway of steroidal A,B,C,D-ring breakdown by *C. testosteroni* TA441.

**IMPORTANCE:** Studies on bacterial steroid degradation were initiated more than 50 years ago, primarily to obtain materials for steroid drugs. Now, their implications for the environment and humans, especially in relation to the infection and the brain-gut-microbiota axis, are attracting increasing attention. *Comamonas testosteroni* TA441 is the leading model of bacterial aerobic steroid degradation with the ability to break down cholic acid, the main component of bile acids. Bile acids are known for their variety of physiological activities according as their substituent group(s). In this study, we identified and functionally characterized the genes for removal of C12 hydroxyl groups and provide a comprehensive summary of the entire A,B,C,D-ring degradation pathway by *C. testosteroni* TA441 as the representable bacterial aerobic degradation process of the steroid core structure.

## INTRODUCTION

Several bacteria, especially *Rhodococcus equi* and *Comamonas testosteroni,* are known for their ability to degrade steroid compounds. The underlying mechanism has been studied extensively to obtain materials for the synthesis of steroidal drugs in the 1960s (1–5). These studies led to the identification of the main intermediates in the A- and B-ring degradation processes.

Nowadays, steroid compounds and the bacteria that metabolize them are attracting attention for their impact on human health, especially in relation to the infection and the brain-gut-microbiota axis. The *mce4* operon, which encodes a cholesterol import system, is essential for persistence of *Mycobacterium tuberculosis* in the lungs of chronically infected animals and for growth within interferon-gamma-activated macrophages (6). Cholesterol catabolism and its utilization by *M. tuberculosis* are important for the pathogen maintenance in the host (7). The primary bile acid, which is synthetized from cholesterol and secreted into the intestine, is converted to secondary bile acid and other cholic acid derivatives by gut bacteria. These acids affect the gut flora and human health. Gut flora is recently attracting increasing attention for the influence on the brain-gut axis, being called the brain-gut-microbiota axis (8). *Comamonas testosteroni* is an environmental bacterium unable to degrade cholesterol, but capable of utilizing cholic acid as a carbon and energy source, whereas some *Comamonas* species are emerging as important opportunistic pathogens (9).

Genetic studies on aerobic steroid degradation by *C. testosteroni*, now the leading bacterial model of this process, started around the year 1990. Thereafter, the enzymes catalyzing the early steps of steroidal degradation, 17β-dehydrogenase (10–13), 3α-dehydrogenase (14–18), 3-oxo-Δ5-steroid isomerase (19, 20), Δ1-dehydrogenase (21), Δ4-dehydrogenase (22), and the positive regulator (23), were identified. However, degradation of steroidal A,B,C,D-rings had remained unclear until we revealed the steroid degradation process in *C. testosteroni* TA441. TA441 degrades steroids through aromatization of the A-ring, along with cleavage of the B-ring, and subsequent cleavage and degradation of the aromatized A-ring. A similar process has been reported for other bacteria, including *Pseudomonas*, *Sphingobium*, *Azoarcus*, and *Rhodococcus* (24).

Degradation of the C,D-ring and cleaved B-ring occurs via β-oxidation (Fig. 1) and a similar process has been reported for *M. tuberculosis* (25). According to the paper on the degradation of the C,D-ring and cleaved B-ring in *M. tuberculosis* (25), *C. testosteroni* CNB-2 also has the similar process. However, to the best of our knowledge, steroid degradation has not been reported for CNB-2. The genomic analysis of CNB-2 showed that it contains putative steroid-degrading genes that are almost identical to those of TA441 (26). Genes responsible for aromatic ring degradation and those mediating β-oxidation respectively form a cluster, which are located at the two ends of the 120-kb mega-cluster of steroid-degrading genes of TA441 (Fig. 1). The function of most genes in these two clusters has been elucidated, except for ORF6, ORF7, ORF25, and ORF26. The latter two are not required for steroid degradation, leaving only ORF6 and ORF7 to be characterized. Here, we describe the role of ORF6 and ORF7 in steroid degradation, and provide a comprehensive summary of the processes guiding the degradation of steroidal A,B,C,D-rings in *C. testosteroni* TA441.

**Fig 1.**
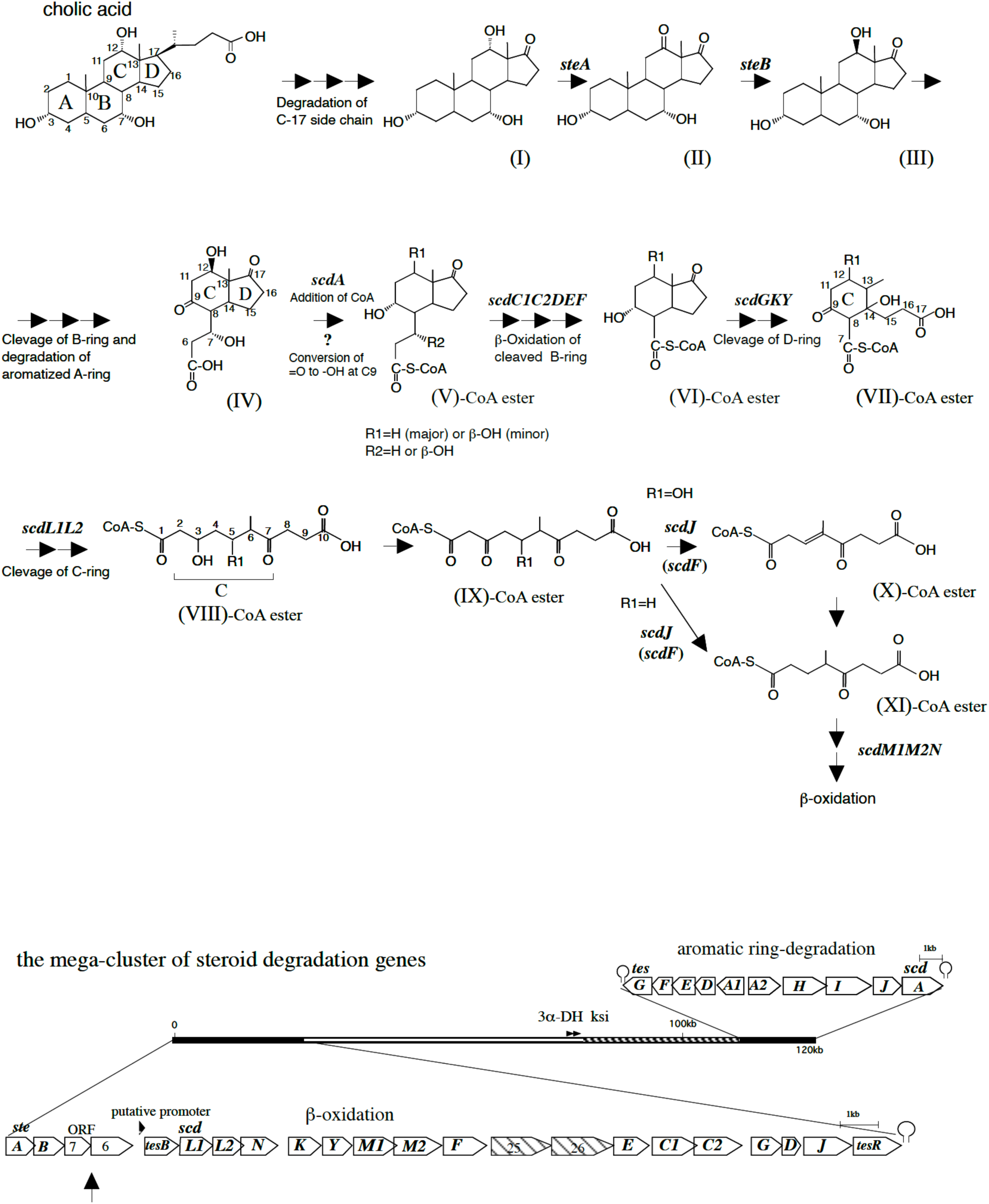
Abbreviated steroid degradation pathway of *Comamonas testosteroni* TA441 indicating reactions and compounds described in this study. The mega-cluster of steroid degradation genes in *C. testosteroni* TA441 is shown below the pathway; the aromatic ring-degradation gene cluster (*tesG* to *scdA*) and the β-oxidation gene cluster (*tesB* to *tesR*) locate both ends of this 120kb-mega cluster. ORF7 and ORF6, indicated with an arrow and in the DNA region upstream of the β-oxidation gene cluster, are left unidentified. Compounds are; 3α,7α,12α-trihydroxy-androstan-17-one, (**I**); 3α,7α-dihydroxy-androstane-12,17-dione, (**II**); 3α,7α,12β-trihydroxy-androstan-17-one, (**III**); 7α,12β-dihydroxy-9,17-dioxo-1,2,3,4,10,19-hexanorandrostan-5-oic acid, (**IV**); 9α-hydroxy-17-oxo-1,2,3,4,10,19-hexanorandrostan-5-oic acid (R1, R2=H), (**V**); 9α-hydroxy-17-oxo-1,2,3,4,5,6,10,19-octanorandrostan-7-oic acid (R1=H) (**VI**); 14-hydroxy-9-oxo-1,2,3,4,5,6,10,19-octanor-13,17-secoandrostane-7,17-dioic acid (R1=H), (**VII**); 6-methyl-3,7-dioxo-decane-1,10-dioic acid-CoA ester (R1=H), (**VIII**); 3-hydroxy-6-methyl-7-oxo-decane-1,10-dioic acid (R1=H), (**IX**); 4-methyl-5-oxo-oct-3-ene-1,8-dioic acid (**X**), and 4-methyl-5-oxo-octane-1,8-dioic acid (**XI**). 3α-DH: 3α-Hydroxy-dehydrogenase gene. ksi: 3-ketosteroid Δ4-5 isomerase gene.

## RESULTS AND DISCUSSION

### Identification of compounds in the culture of an ORF6-disrupted mutant incubated with cholic acid

*C. testosterone* TA441 degrades steroidal A,B,C, and D-rings with the pathway similar to aromatic compound degradation for A,B-ring cleavage followed by C,D, and cleaved B-ring degradation mainly by β-oxidation. The degradation genes are encoded in two clusters on both end of the 120kb-mega cluster of steroid degradation genes; one consists of those for aromatic compound degradation and the other for β-oxidation (Fig. 1). ORF6 and ORF7 are the only two genes in these clusters whose function remains unknown. They are in the DNA region just upstream of the β-oxidation gene cluster, and downstream of the genes encoding SteA and SteB, which catalyze the conversion of the C12α hydroxyl to the C12β hydroxyl via a ketone group (Fig. 1) (27). Previously, *steAB*, ORF7, and ORF6 were shown to be involved in cholic acid degradation; while their disruption had no influence on growth on testosterone (27). A homology search revealed that the ORF7-encoded enzyme belonged to the nuclear transport factor 2 family, which includes Δ5-3-ketosteroid isomerases and LinA. LinA is a dehalogenase with hydrolase activity, which converts γ-hexachlorocyclohexane to γ -pentachlorocyclohexene (28). The ORF6-encoded enzyme was revealed to be an old yellow enzyme-like alkene reductase whose substrate was not clear. Here, gene-disrupted mutants of ORF7 and ORF6, denoted respectively as ORF7^-^ and ORF6^-^, were incubated with cholic acid and its analogs, chenodeoxycholic acid, deoxycholic acid, and lithocholic acid, and the cultures were analyzed by high-performance liquid chromatography (HPLC) with UV detection. Cholic acid harbors hydroxyl groups at positions C7 and C12. Chenodeoxycholic acid contains a hydroxyl group at position C7 and deoxycholic acid at position C12. Lithocholic acid does not have either of them (Fig. 2). Candidate intermediate compounds were detected only in the ORF6^-^ culture incubated with cholic acid (Fig. 2). The three compounds, denoted as 6a, 6b, and 6c (Fig. 2), were isolated and identified by fast atom bombardment mass spectrometry as having mass/charge ratios (*m/z*) 219, 219, and 217 (M+H^+^), respectively. Accordingly, their respective molecular formulae were deduced to be C_13_H_14_O_3_, C_13_H_14_O_3_, and C_13_H_12_O_3_. Based on nuclear magnetic resonance (NMR) analysis, 6a, 6b, and 6c were identified as the lactone forms of 17-hydroxy-9-oxo-1,2,3,4,10,19-hexanorandrost-6,10(12)-dien-5-oic acid (**XII**), 9,17-dioxo-1,2,3,4,10,19-hexanorandrost-6-en-5-oic acid (**XIII**), and 9,17-dioxo-1,2,3,4,10,19-hexanorandrost-6,10(12)-dien-5-oic acid (**XIV**), respectively. The corresponding structures are presented in Fig. 2 and NMR data are listed in Table 1. These compounds had a ketone moiety at position C9 and, therefore, they corresponded to the intermediates found before β-oxidation of the cleaved B-ring (cf. Fig. 1, **IV** to **V**).

**Fig 2.**
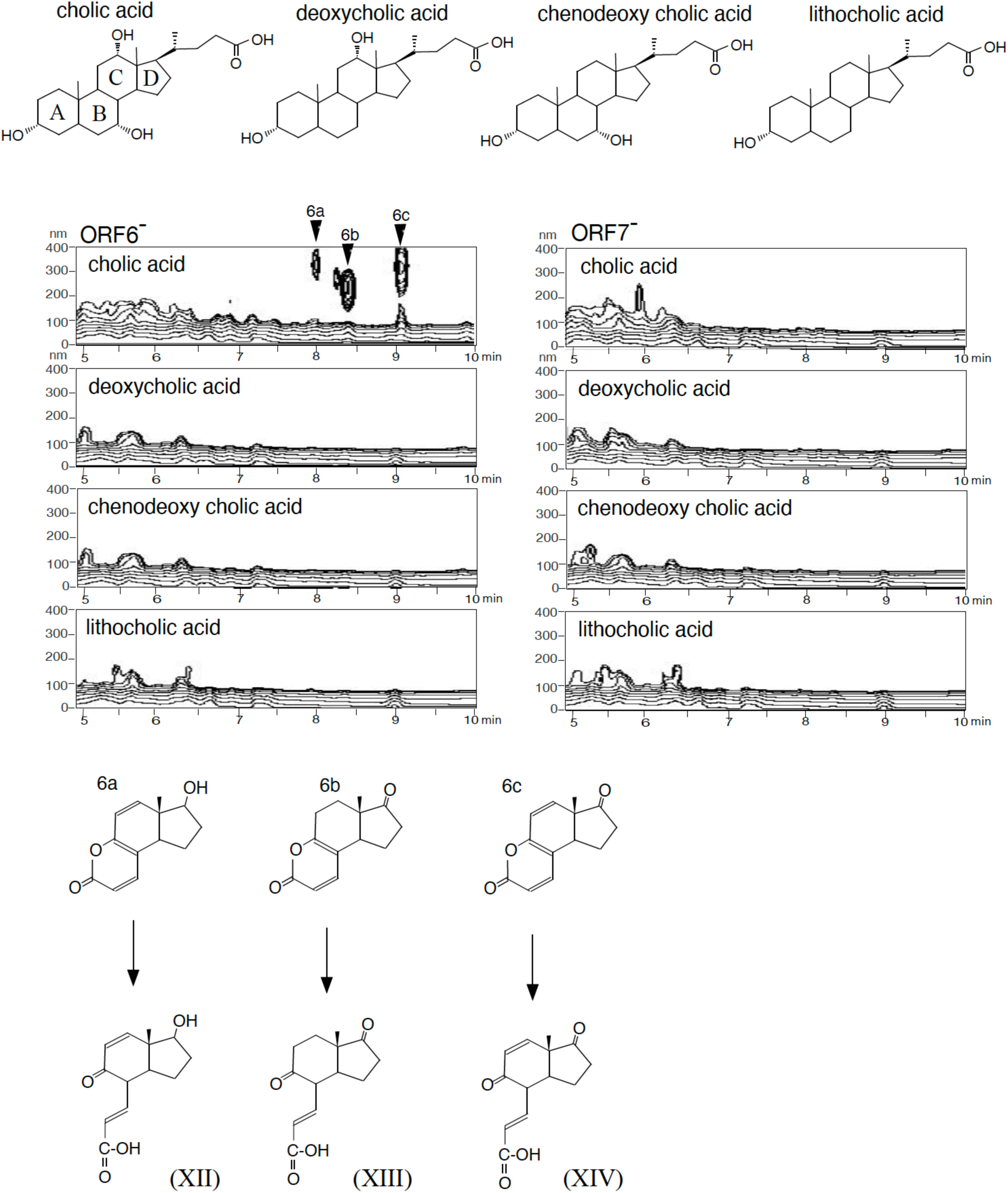
HPLC analysis of the culture of the ORF7^-^ and ORF6^-^ mutants incubated with cholic acid, deoxycholic acid, chenodeoxycholic acid, and lithocholic acid, and the compounds identified from the culture of ORF6^-^ mutant with cholic acid (6a - 6c). These compounds (6a - 6c) were lactones of 17-hydroxy-9-oxo-1,2,3,4,10,19-hexanorandrost-6,10-dien-5-oic acid (**XII**), 9,17-dioxo-1,2,3,4,10,19-hexanorandrost-6-en-5-oic acid (**XIII**), and 9,17-dioxo-1,2,3,4,10,19-hexanorandrost-6,10-dien-5-oic acid (**XIV**), respectively. In HPLC, the vertical axis indicates wavelength (nm), the horizontal axis indicates RT (min), and the UV absorbance of each compound is represented in contour.

**Table 1.** NMR data of compounds accumulated by ORF6-disrupted mutant incubated with cholic acid.

| No | Compound 6a (RT=6.7 min)<br>(in CDCl<sub>3</sub>) |  | Compound 6b (RT=7.6 min)<br>(in CDCl<sub>3</sub>) |  | Compound 6c (RT=8.7 min)<br>(in CDCl<sub>3</sub>) |  |
| --- | --- | --- | --- | --- | --- | --- |
|  | <sup>13</sup>C-NMR<br>[δ (ppm)] | <sup>1</sup>H-NMR<br>[δ (ppm)], J (Hz)] | <sup>13</sup>C-NMR<br>[δ (ppm)] | <sup>1</sup>H-NMR<br>[δ (ppm)], J (Hz)] | <sup>13</sup>C-NMR<br>[δ (ppm)] | <sup>1</sup>H-NMR<br>[δ (ppm)], J (Hz)] |
| 5 | 164.60 |  | 164.81 |  | 164.30 |  |
| 6 | 113.44 | 6.17 d (9.2) | 113.66 | 6.20 d (9.2) | 114.30 | 6.26 d (4.1) |
| 7 | 145.13 | 7.41 d (9.2) | 145.35 | 7.45 d (9.2) | 144.46 | 7.56 d (9.2) |
| 8 | 116.12 |  | 115.20 |  | 115.25 |  |
| 9 | 158.39 |  | 161.94 |  | 158.45 |  |
| 11 | 121.60 | 6.14 d (9.8) | 26.40 | 2.75 m | 122.69 | 6.24 d (4.6) |
| 12 | 147.05 | 6.67 d (9.8) | 28.18 | 1.72 ddd (13.3, 9.4, 9.2) | 141.07 | 6.66 d (10.1) |
| 13 | 47.92 |  | 49.00 |  | 51.10 |  |
| 14 | 43.48 | 2.83 (7.5, 5.1) | 44.00 | 2.89 | 44.03 | 3.14 (13.1, 6.2) |
| 15 | 21.07 | 1.94 m | 21.51 | 1.85 m | 19.43 | 2.06 m |
|  |  | 1.80 dddd(12.0, 12.0, 12.0, 6.0) |  | 2.34 m |  | 2.30 m |
| 16 | 31.71 | 2.30 m | 37.14 | 2.32 m | 37.41 | 2.46 ddd (19.0, 9.5, 9.5) |
|  |  | 1.71 m |  | 2.65 m |  | 2.68 dd (16.7, 8.7) |
| 17 | 75.68 | 4.09 dd (9.2, 7.8) | 219.43 |  | 214.12 |  |
| 18 | 10.55 | 0.78 s | 13.16 | 0.87 s | 13.22 | 0.89 s |

Abbreviations for NMR signals are as follows s, singlet; d, doublet; m, multiplet.

### Analysis of the culture of ORF7^-^ and ORF6^-^ mutants incubated with cholic acid and its analogs

Because HPLC could not detect compounds with weak UV absorption, ORF7^-^ and ORF6^-^ mutants were incubated individually with cholic acid, deoxycholic acid, chenodeoxycholic acid, and lithocholic acid, and the respective cultures were then analyzed by reverse-phase liquid chromatography with tandem mass spectrometry (LC-MS/MS). A compound with *m/z* 253 at retention time (RT) = 4.5 min and another with *m/z* 255 at RT = 4.3 min were detected in large amounts in the ORF7^-^ culture with cholic acid or deoxycholic acid (Fig. 3A1,2 and Fig. 3C1,2). Somewhat lower but nevertheless substantial amounts of the same compound were detected in the ORF6^-^ culture (Fig. 3B1,2 and Fig. 3D1,2). These compounds were not detected in cultures incubated with chenodeoxycholic acid or lithocholic acid, which do not have a hydroxyl group at position C12 (Fig. 3A3,4; Fig. 3B3,4; Fig. 3C3,4; and Fig. 3D3,4). Mass chromatograms of *m/z* 269, 271, 237, 235, 211, 227, 181, 197, 241, 259, 245, 243, and 201 of the ORF6^-^ culture with cholic acid (Fig. S1-1), ORF7^-^ culture with cholic acid (Fig. S1-2), ORF7^-^ culture with deoxycholic acid (Fig. S1-3), and ORF7^-^ culture with chenodeoxycholic acid (Fig. S1-4) with the mass spectra (Fig. S2) are presented in Supplemental Material. Based on the peak with *m/z* 253 at RT = 4.5 min and its major fragments with *m/z* 253, 235, 191, 163, and 149 (Fig. S2A1), as well as the peak with *m/z* 255 at RT = 4.3 min and its major fragments with *m/z* 255, 237, and 193 (Fig. S2B1), the two peaks were assigned to 12β-hydroxy-9,17-dioxo-1,2,3,4,10,19-hexanorandrostan-5-oic acid (**XV**) and 12β,17-dihydroxy-9-oxo-1,2,3,4,10,19-hexanorandrostan-5-oic acid (**XVI**), respectively. Other characteristic peaks were observed in the mass chromatograms of *m/z* 235 (Fig. 3E1,2), *m/z* 227 (Fig. 3E3,4), and *m/z* 241 (Fig. 3E5,6).

**Fig 3.**
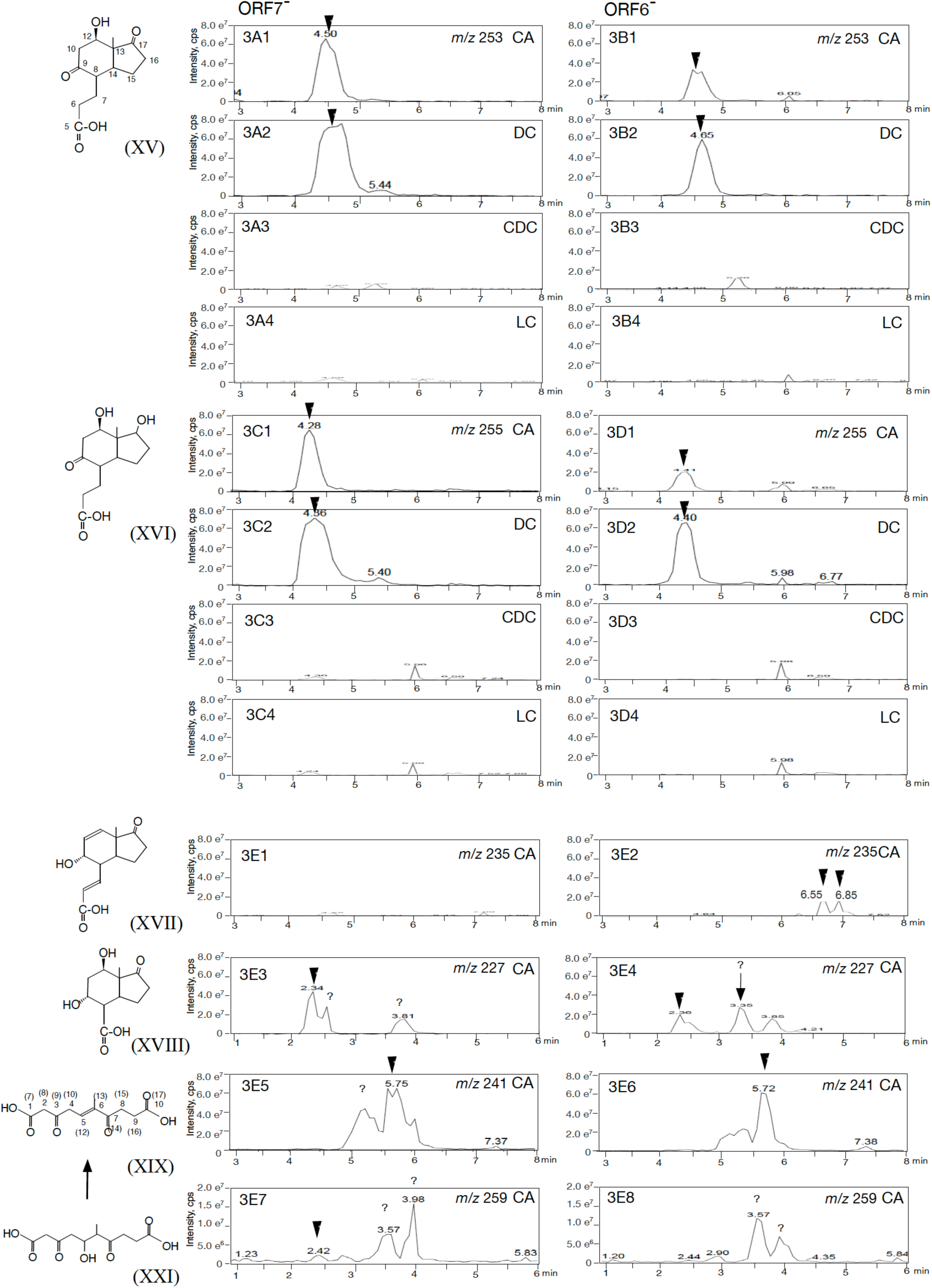
LC/MS/MS analysis of the culture of the ORF7^-^ and ORF6^-^ mutants incubated with cholic acid (CA), deoxycholic acid (DC), chenodeoxycholic acid (CDC), and lithocholic acid (LC). Chromatograms of *m/z* 253 (3A1-4, 3B1-4), *m/z* 255 (3C1-4, 3D1-4), *m/z* 235 (3E1, 3E2), *m/z* 227 (3E3, 3E4), *m/z* 241 (3E5, 3E6), and *m/z* 259 (3E7, 3E8) are shown. Two peaks with *m/z* of 241 at around RT = 5.7 min in ORF7^-^ culture (3E5) showed the almost identical mass spectrums (Fig. S2J4 in Supplemental Material). Closed arrows indicate the peak of the compound shown on the left side of the columns. For **XIX** and **XXI**, the carbons are given new numbers according to IUPAC numbering. Numbers in parentheses on **XIX** are based on the steroidal numbering. Compounds are; 12β-hydroxy-9,17-dioxo-1,2,3,4,10,19-hexanorandrostan-5-oic acid (**XV**), 12β,17-dihydroxy-9-oxo-1,2,3,4,10,19-hexanorandrostan-5-oic acid (**XVI**), 9α-hydroxy-17-oxo-1,2,3,4,10,19-hexanorandrosta-6,10(12)-dien-5-oic acid (**XVII**), 9α,12β-dihydroxy-17-oxo-1,2,3,4,5,6,10,19-octanorandrostan-7-oic acid (**XVIII**), 6-methyl-3,7-dioxo-dec-5-ene-1,10-dioic acid (**XIX**), and 5-hydroxy-6-methyl-3,7-dioxo-decane-1,10-dioic acid (**XXI**). The vertical axis indicates intensity (count/sec) and the horizontal axis indicates RT (min). Mass chromatograms of *m/z* 269, 271, 237, 235, 211, 227, 181, 197, 241, 259, 245, 243, and 201 are presented in Supplemental Material Fig. S1-1 to S1-4. Fig. S1-1 shows ORF6^-^ culture with cholic acid. Fig. S1-2, S1-3, and S1-4 show ORF7^-^ culture with cholic acid, deoxycholic acid, and chenodeoxycholic acid, respectively.

9α-Hydroxy-17-oxo-1,2,3,4,10,19-hexanorandrosta-6,10(12)-dien-5-oic acid (**XVII**), a compound with a C10(12) double bond, was detected as peaks with *m/z* 235 at RT = 6.55 and 6.85 min in the ORF6^-^ culture (Fig. 3E1), while it was absent in the ORF7^-^ culture (Fig. 3E2 and Fig. S1F3,5 in Supplemental Material). In contrast, the peak with *m/z* 227 at RT = 2.3 min showed accumulation in the ORF7^-^ culture (Fig. 3E3) but lower levels in the ORF6^-^ culture (Fig. 3E4). This compound presented major fragments with *m/z* 227, 209, 183, and 155 and was identified as 9α,12β-dihydroxy-17-oxo-1,2,3,4,5,6,10,19-octanorandrostan-7-oic acid (**XVIII**) (Fig. S2G1 in Supplemental Material). All C-ring-containing intermediates detected in ORF7^-^ cultures incubated with cholic acid or deoxycholic acid harbored a hydroxyl group at C12 (Fig. 3 and Fig. S1-2,3 in Supplemental Material). Based on these findings and homology search result, the ORF7-encoded enzyme was presumed to be the dehydrase for the C12β hydroxyl group. ScdA is a coenzyme A (CoA)-transferase, which adds CoA to the C5 carboxylic group, thereby initiating β-oxidation. The main compound found to accumulate in an ScdA^-^ culture with cholic acid was 7α,12β-dihydroxy-9,17-dioxo-1,2,3,4,10,19-hexanorandrostan-5-oic acid (29). At the same time, none of the intermediates during β-oxidation of the cleaved B-ring identified in the previous studies contained the C12β hydroxyl group, indicating that the hydroxyl group is basically removed prior to β-oxidation. Present data indicate that C,D-ring cleavage proceeds less efficiently in the presence than in the absence of the C12β hydroxyl group.

The mass spectrum of the large peak with *m/z* 241 detected at RT = 5.72 min was identical to that of 6-methyl-3,7-dioxo-dec-5-ene-1,10-dioic acid (**XIX**). Note that we used IUPAC numbering for compounds without steroidal rings, and steroidal numbering for those with at least one steroidal ring. This compound was initially identified as an intermediate produced after C-ring cleavage by ScdL1L2 in cultures incubated with steroids harboring a hydroxyl group at C12 (accompanying paper). **XIX** and 3-hydroxy-6-methyl-7-oxo-dec-5-ene-1,10-dioic acid (**XX**) (Fig. S1-1N,2N,3N in Supplemental Material), whose main fragments possessed *m/z* values of 243, 199, 155, 153, 111, and 109 (Fig. S2M in Supplemental Material), were the only intermediate compounds with a double bond at C5 (C12 of the C-ring), and which accumulated more in the ORF7^-^ than in the ORF6^-^ culture (Fig. 3E5,6, and Fig. S1-2N,1N in Supplemental Material). Therefore, the compounds originally present in the culture were hypothesized as 5-hydroxy-6-methyl-3,7-dioxo-decane-1,10-dioic acid (**XXI**) (Fig. 3E7) and 3,5-dihydroxy-6-methyl-7-oxo-decane-1,10-dioic acid (**XXII**) (Fig. S1-1N,2N,3N in Supplemental Material), respectively. They were thought to become dehydrated during extraction and analysis in acidic conditions.

In ORF6^-^ cultures incubated with cholic acid and deoxycholic acid, compounds with a double bond at C10(12) and a single bond at C10(12) accompanied those with a C12 hydroxyl group (Fig. 3 and Fig. S1 in Supplemental Material). Intermediates harboring the C12 hydroxyl group were less abundant in ORF6^-^ cultures than in ORF7^-^ cultures, suggesting that the reaction catalyzed by the ORF6-encoded enzyme occurred after dehydration by the ORF7-encoded enzyme. Compounds with a double bond at C10(12) in the C-ring, including **XII**, **XIV**, **XVII**, and 9,17-dioxo-1,2,3,4,10,19-hexanorandrost-10(12)-en-5-oic acid (**XXIII**), were detected only in the ORF6^-^ culture (Fig. 3E2 and Fig. S1-1E in Supplemental Material). Two peaks with *m/z* 235 at RT = 6.55 and 6.85 min showed almost identical mass spectra, characterized by major fragments with *m/z* 235, 217, 191, 173, 163, 135, 123, and 95 (Fig. S2E3,5 in Supplemental Material), and were thought to be stereoisomers. The above data, together with homology search results, pointed to the ORF6-encoded enzyme being a reductase, which converted a double bond to a single bond at C10(12) in the C-ring. The presence of compounds with a single bond at C10(12) in ORF6^-^ cultures implied the existence of at least another reductase capable of acting on the double bond at C10(12). A peak with *m/z* 227 at RT = 3.3 min was detected exclusively in the ORF6^-^ culture incubated with cholic acid or deoxycholic acid (Fig. 3E4), and was thought to be one of the key intermediates that could confirm the function of the ORF6-encoded enzyme. Unfortunately, we were unable to deduce the compound’s structure from the mass spectrum, which was characterized by a large fragment with *m/z* 227 and minor fragments with *m/z* 209, 183, 165, and 155 (Fig. S2G2 in Supplemental Material).

### Complementation of enzymes encoded by ORF7 and ORF6

To further investigate the function of ORF7- and ORF6-encoded enzymes, different combinations of mutant strains and plasmids were generated. These included ORF7^-^ carrying the broad-host-range plasmid pMFY42 (negative control), ORF7^-^ carrying a pMFY42-based plasmid encoding ORF7 (pMFYORF7), ORF6^-^ carrying pMFY42 (negative control), ORF6^-^ carrying a pMFY42-based plasmid encoding ORF6 (pMFYORF6), double ORF7^-^ORF6^-^ mutant (ORF7^-^6^-^) carrying pMFY42 (negative control), ORF7^-^6^-^ carrying pMFYORF7, and ORF7^-^6^-^ carrying a pMFY42-based plasmid encoding ORF7 and ORF6 (pMFYORF76) (Tables 2–4). The mutants were cultivated with cholic acid for 7 days and the cultures were analyzed by LC-MS/MS.

**Table 2.** strains

| strains | Characteristics | Source or reference |
| --- | --- | --- |
| TA441 | Wild-type | [45] |
| ORF6<sup>-</sup> | ORF6: :Km<sup>r</sup> mutant (*Bgl*III site) of TA441 | [27] |
| ORF7<sup>-</sup> | ORF7: :Km<sup>r</sup> mutant (*Apa*I site) of TA441 | [27] |
| ORF7,6<sup>-</sup> | ORF7,6: :Km<sup>r</sup> mutant (*Apa*I site to *Bgl*III site ) of TA441 | this work |

Km<sup>r</sup> : Km-resistance, [45] Arai et al. 1998. Microbiology **144**: 2895-2903 [27] Horinouchi et al. 2008. J Bacteriol **190**:5545-5554

**Table 3.** plasmids

| plasmids | Characteristics | Source or reference |
| --- | --- | --- |
| pUC19 | Ap<sup>r</sup>, *lacZ* | [44] |
| pMFY42 | Tc<sup>r</sup>, Km<sup>r</sup>, RSF1010-based broad host range plasmid | [28] |
| pMFYORF6 | pMFY42 derivative carrying ORF6 | this work |
| pMFYORF7 | pMFY42 derivative carrying ORF7 | this work |
| pMFYORF76 | pMFY42 derivative carrying ORF7,6 | this work |
| pMFYMhpRA | pUC19 derivative carrying *mhpR* and the promoter of *mhp* genes | [*] |
| pMFYMhpORF76 | pMFYMhpRA derivative carrying ORF7,6 | this work |

Km<sup>r</sup> : Km-resistance [44] Vieira, J., and Messing, J. 1987. Methods Enzymol. **153**: 3-11. [28] Nagata Y. et al. 1993. J Bacteriol **175**:6403-6410 [\*] Horinouchi M, et al. submitted.

**Table 4.**
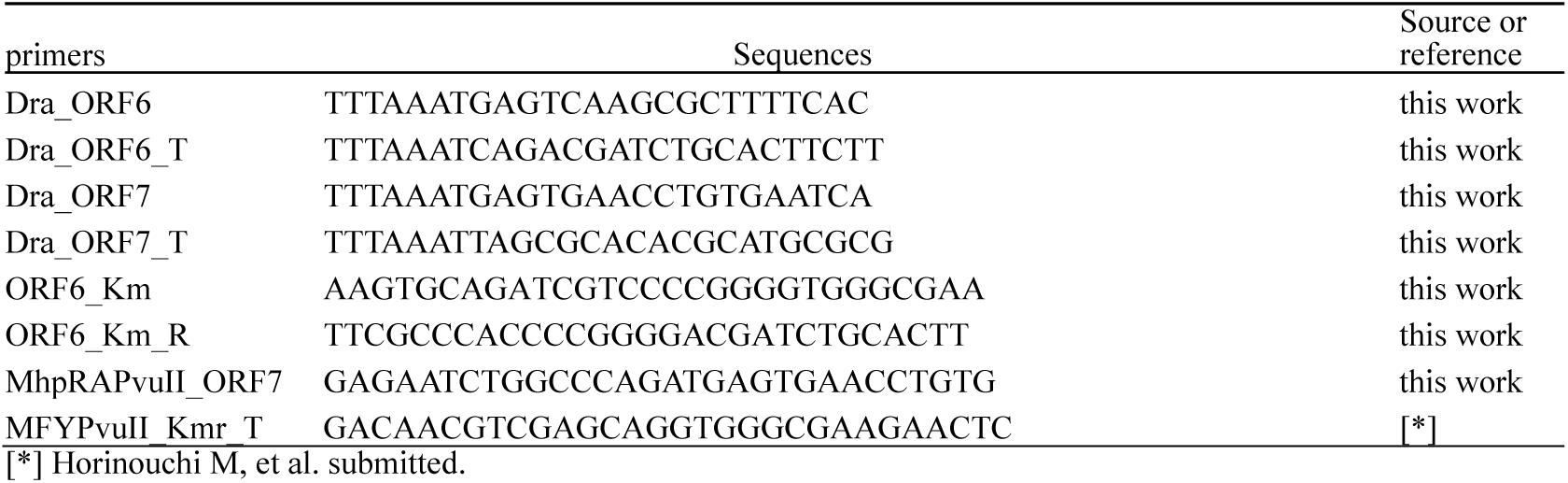
primers

The mass chromatograms of *m/z* 253, 235, and 237 are shown in Fig. 4 and the mass chromatograms of *m/z* 227, 241, 197, and 201 are shown in Fig. S3 in Supplemental Material. **XV** was less abundant in all complemented mutants, ORF7^-^ carrying pMFYORF7 (Fig. 4-1B), ORF6^-^ carrying pMFYORF6 (Fig. 4-1D), and ORF7^-^6^-^ carrying pMFYORF76 (Fig. 4-1H). A similar, albeit less pronounced drop was observed for **XV** in ORF7^-^6^-^ carrying pMFYORF7 (Fig. 4-1G). **XVII** and **XXIII**, both of which have a double bond at C10(12), were detected in the culture of ORF7^-^ carrying pMFYORF7 (Fig. 4-2B), ORF6^-^ carrying pMFY42 (Fig. 4-2C), ORF6^-^ carrying pMFYORF6 (Fig. 4-2D), and ORF7^-^6^-^ carrying pMFYORF7 (Fig. 4-2G). Slightly lower levels of **XVII** and **XXIII** were detected in the culture of ORF6^-^ carrying pMFYORF6 compared to the one with ORF6^-^ carrying pMFY42. **XIII**, a compound with a single bond at C10(12), was detected in the cultures of the three complemented mutants (Fig. 4-2B,D,H). The data for the ORF7-culture incubated with chenodeoxycholic acid, which does not have a C12 hydroxyl group, are presented in Fig. 4-2E as an authentic for **XIII** . These results confirmed the ability of the ORF7-encoded enzyme to dehydrate the C12 hydroxyl group of **XV**, thereby generating **XVII** and **XXIII**. 9,17-Dioxo-1,2,3,4,10,19-hexanorandrostan-5-oic acid, (3aα-*H*-4α [3′-propionic acid]-7aβ-methylhexahydro-1,5-indanedione) (R1, R2 = H) (**XXIV**), which has a single bond at C10(12), was detected in the culture of ORF7^-^ carrying pMFYORF7 (Fig. 4-3B), ORF6^-^ carrying pMFY42 (Fig. 4-3C), ORF6-carrying pMFYORF6 (Fig. 4-3D), and ORF7^-^6^-^ carrying pMFYORF7 (Fig. 4-3G). The corresponding mass spectra are presented in Fig. S2D3 in Supplemental Material. It is difficult to distinguish **XXIV** from 9α-hydroxy-17-oxo-1,2,3,4,10,19-hexanorandrost-6-en-5-oic acid (R1 = H) (**XXV**) based on the mass chromatogram, but both can be regarded as **XV** derivatives, with a single bond at C10(12) and without a hydroxyl group at C12. Less **XV** was detected in the culture of ORF7^-^6^-^ carrying pMFYORF7 (Fig. 4-1G) than in the corresponding control carrying pMFY42 (Fig. 4-1F), although this reduction was less pronounced than the one observed with ORF7^-^ carrying pMFYORF7 (Fig. 4-1B). **XV** dropped to nearly undetectable levels in the culture of ORF7^-^6^-^ carrying pMFYORF76 (Fig. 4-1H). These results implied that the C12β hydroxyl group was removed efficiently when both ORF7- and ORF6-encoded enzymes were expressed together.

**Fig 4.**
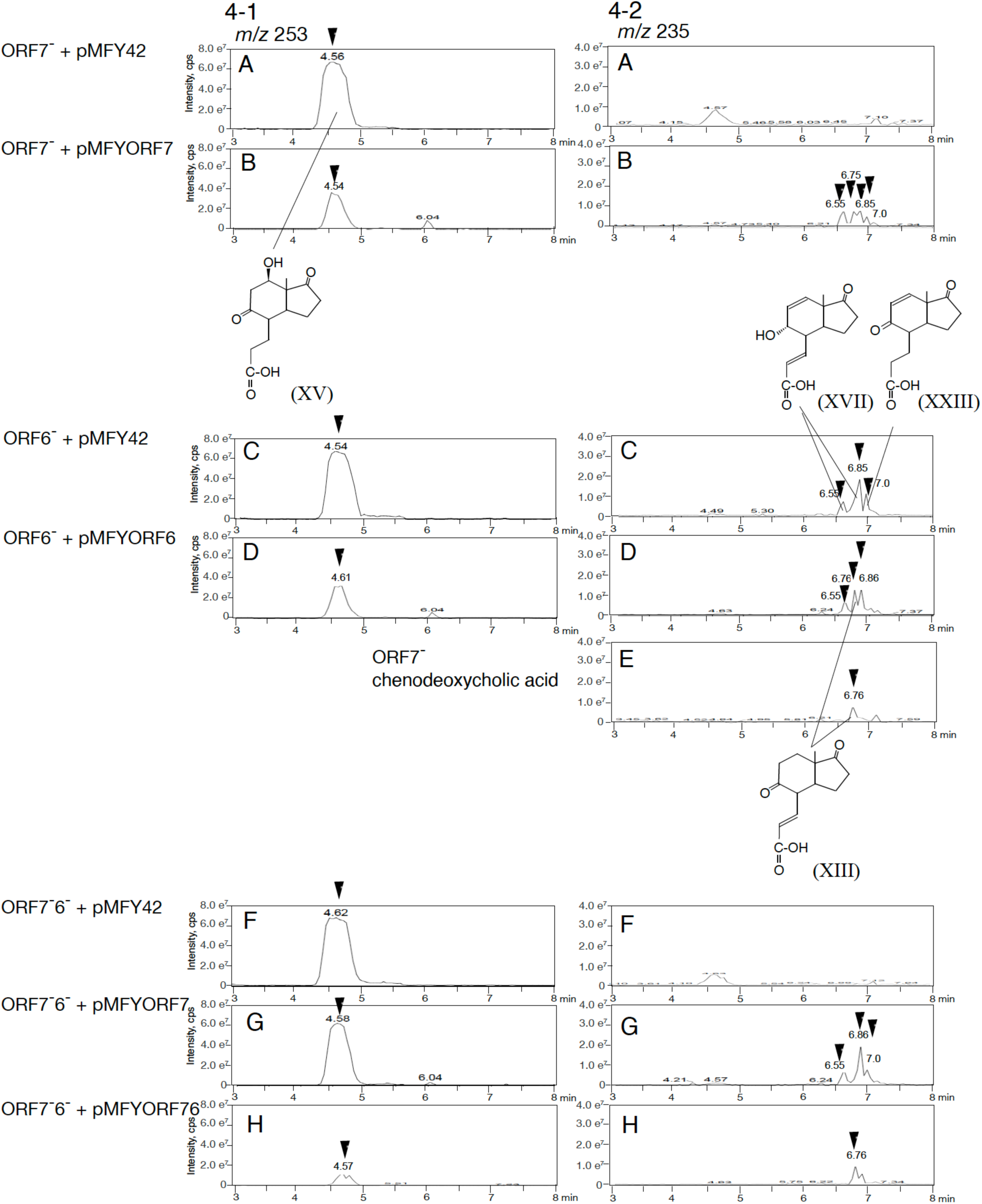
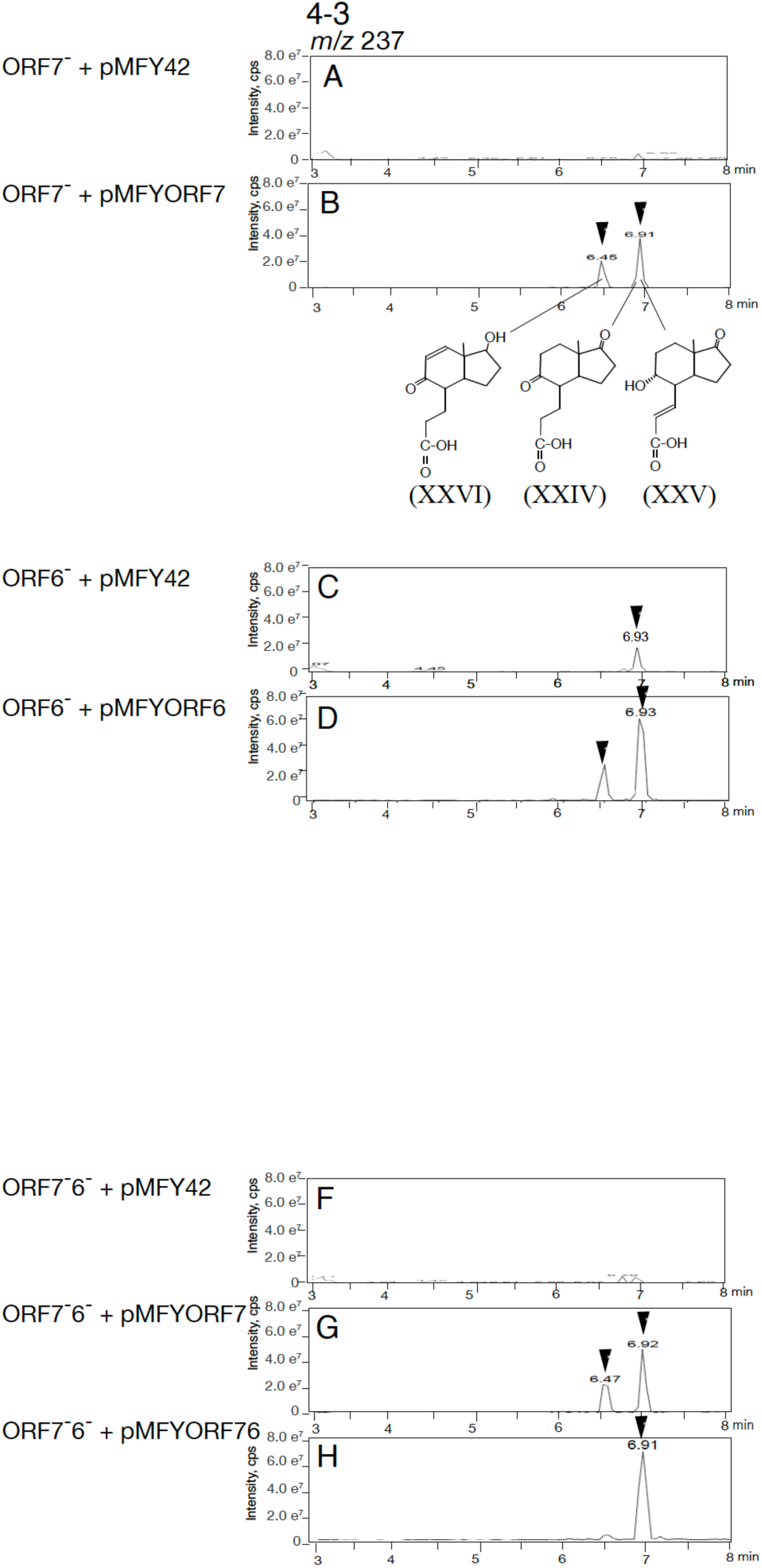
Complementation experiments with ORF7^-^, ORF6^-^, and ORF7^-^6^-^ mutants. The mass chromatograms of each mutant culture incubated with 0.1% cholic acid for 7days are shown. Mutants are; ORF7^-^ mutant carrying pMFY42 (ORF7^-^ with pMFY42) (negative control) (A), carrying pMFYORF7 (pMFY42 carrying ORF7) (B), ORF6^-^ mutant carrying pMFY42 (ORF6^-^ with pMFY42) (negative control) (C), carrying pMFYORF6 (ORF6^-^ with pMFYORF6) (D), ORF7^-^ mutant incubated with chenodeoxycholic acid (as an authentic for **XIII**) (E), ORF7^-^6^-^ mutant carrying pMFY42 (ORF7^-^6^-^ with pMFY42) (negative control) (F), carrying pMFYORF7 (G), and carrying pMFYORF76 (H). Mass chromatograms of *m/z* 253 (4-1), *m/z* 235 (4-2), and *m/z* 237 (4-3) are shown. Mass chromatograms of *m/z* 227, *m/z* 241, *m/z* 197, and *m/z* 201 are shown Fig. S3 in Supplemental Material. Compounds are; 12β-hydroxy-9,17-dioxo-1,2,3,4,10,19-hexanorandrostan-5-oic acid (**XV**), 9α-hydroxy-17-oxo-1,2,3,4,10,19-hexanorandrosta-6,10(12)-dien-5-oic acid (**XVII**), 9α,17-dioxo-1,2,3,4,10,19-hexanorandrost-10(12)-en-5-oic acid (**XXIII**), 9α,17-dioxo-1,2,3,4,10,19-hexanorandrost-6-en-5-oic acid (**XIII**), 9,17-dioxo-1,2,3,4,10,19-hexanorandrostan-5-oic acid (**XXIV**) (R1, R2=H), 9α-hydroxy-17-oxo-1,2,3,4,10,19-hexanorandrost-6-en-5-oic acid (**XXV**), and 17-hydroxy-9-oxo-1,2,3,4,10,19-hexanorandrost-10(12)-en-5-oic acid (**XXVI**). The vertical axis indicates intensity (count/sec) and the horizontal axis indicates RT (min).

### Expression of ORF7- and ORF6-encoded enzymes in the ORF7^-^6^-^ mutant

To further confirm the function of ORF7- and ORF6-encoded enzymes, the pMFYMhpR plasmid was employed. pMFYMhpR is a pMFY42-based plasmid which harbors *mhpR* encoding the positive regulator of 3-(3-hydroxyphenyl)propionic acid (3HPP) degradation genes (*mhp*) (28) and the promoter region of *mhp* genes in TA441 (Fig. 5 above) (accompanying paper). Genes cloned downstream of the *mhp* promoter are induced upon addition of 3HPP. pMFYMhpRORF76, a pMFYMhpR-derivative carrying ORF7 and ORF6 downstream of the *mhp* promoter, was constructed and introduced into the ORF7^-^6^-^ double mutant. ORF7^-^6^-^ carrying pMFYMhpRORF76 and the corresponding negative control (ORF7^-^6^-^ carrying pMFY42) were incubated with cholic acid for 7 days, after which 3HPP was added and the culture was analyzed every day for 3 days. Mass chromatograms of the cultures before and after incubation with 3HPP are reported in Fig. 5. We also analyzed the culture of ORF7^-^6^-^ carrying pMFYMhpRORF76 incubated for 10 days in the absence of any 3HPP supplementation to exclude the possibility that a prolonged incubation altered the compounds. The mass chromatogram of this culture was almost identical to the one obtained after 7 days (data not shown), thus excluding any effect of prolonged incubation time. Compounds harboring a 12β hydroxyl group, such as **XV** (Fig. 5A), **XVIII** (Fig. 5C), and **XXI** (detected as **XIX** in Fig. 5D), exhibited a large drop in amount; whereas **XII, XVII,**

**Fig 5.**
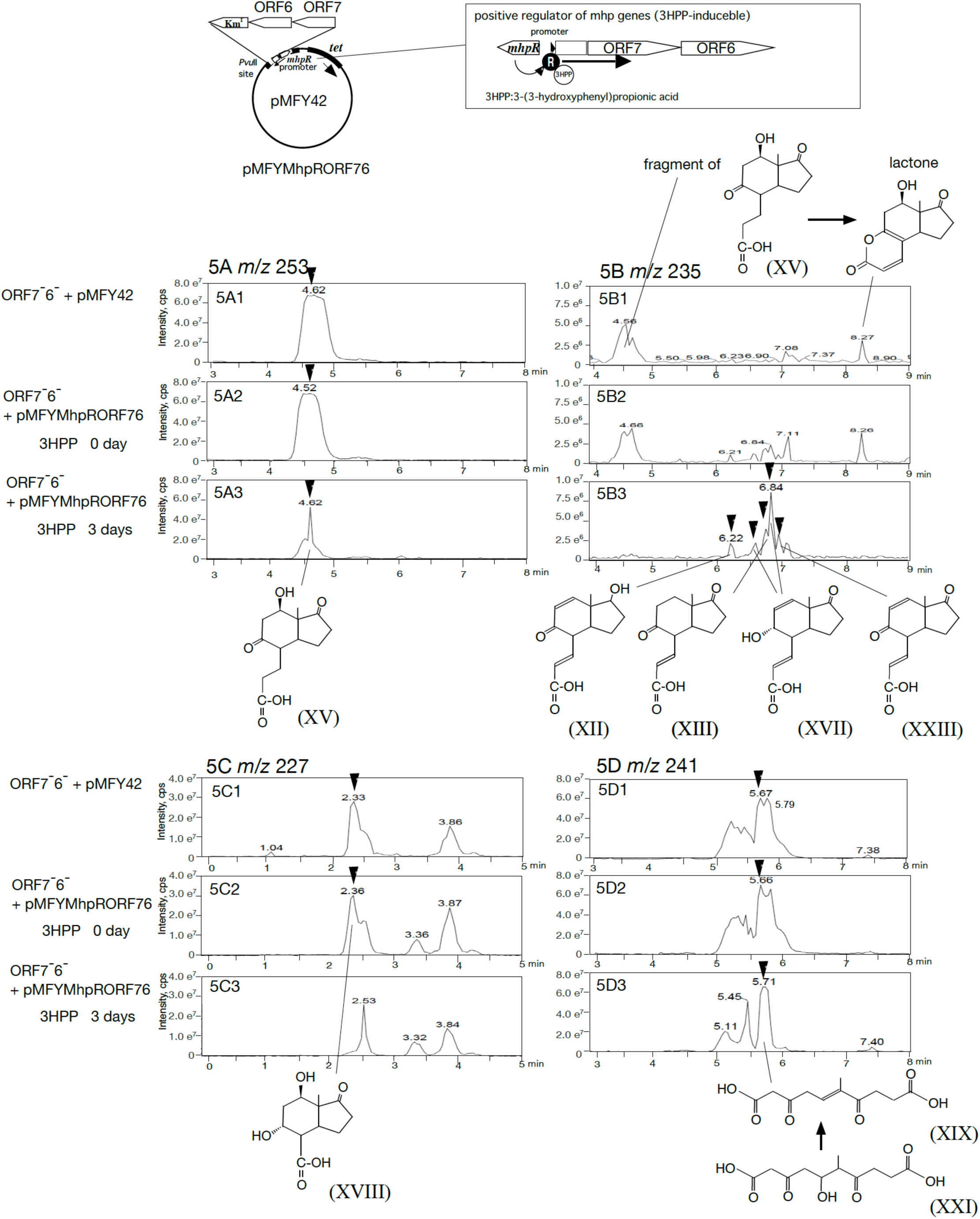
Induction of ORF7 and ORF6 in ORF7^-^6^-^ mutants carrying pMFYMhpRORF76. pMFYMhpRORF76 is a pMFY42 based plasmid carrying positive regulator and the promoter region of 3-(3-hydroxyphenyl)propionic acid (3HPP) degradation genes, in which ORF7 and ORF6 are induced with addition of 3HPP (figure above) (accompanying paper). The mass chromatograms of the culture of ORF7^-^6^-^ with pMFY42 (negative control) (5A1 - 5D1) and ORF7^-^6^-^ with MhpRORF76 (5A2 - 5D2) incubated with 0.1% cholic acid for 7 days, and ORF7^-^6^-^ with MhpRORF76 incubated with 0.1% cholic acid for 7 days followed by additional 3-day incubation with 3HPP (5A3 to 5D3) are shown. Closed arrowheads indicate compounds involved in ORF7- and ORF6-encoded enzymes’ reactions. Compounds are; 12β-hydroxy-9,17-dioxo-1,2,3,4,10,19-hexanorandrostan-5-oic acid (**XV**) (**XXIV**, R1=OH, R2=H), 17-hydroxy-9-oxo-1,2,3,4,10,19-hexanorandrost-10(12)-en-5-oic acid (**XII**), 9α,17-dioxo-1,2,3,4,10,19-hexanorandrost-6-en-5-oic acid (**XIII**), 9α-hydroxy-17-oxo-1,2,3,4,10,19-hexanorandrosta-6,10(12)-dien-5-oic acid (**XVII**), 9α,17-dioxo-1,2,3,4,10,19-hexanorandrost-10(12)-en-5-oic acid (**XXIII**), 9α,12β-dihydroxy-17-oxo-1,2,3,4,5,6,10,19-octanorandrostan-7-oic acid (**XVIII**), 6-methyl-3,7-dioxo-dec-5-ene-1,10-dioic acid (**XIX**), and 5-hydroxy-6-methyl-3,7-dioxo-decane-1,10-dioic (**XXI**). Mass spectrums of these compounds are presented in Fig. S2 in Supplementary Material. The vertical axis indicates intensity (count/sec) and the horizontal axis indicates RT (min).

**XXIII** (compounds with a double bond at C10(12)), and **XIII** (compound with a single bond at C10(12)) (Fig. 5B) exhibited an increase in 3HPP-induced ORF7^-^6^-^ carrying pMFYMhpRORF76. These results confirmed that ORF7- and ORF6-encoded enzymes removed the C12β hydroxyl group via dehydration.

In conclusion, the enzymes encoded by *steA, steB*, ORF7, and ORF6 catalyze the dehydrogenation of the C12α hydroxyl group on the C-ring to a ketone, hydrogenation of the ketone to the C12β hydroxyl group (27), dehydration of the C12β hydroxyl group to produce a double bond at C10(12), and reduction of the double bond to a single bond to remove the C12α hydroxyl group, respectively. Accordingly, ORF7 and ORF6 were named *steC* and *steD*.

### Overall steroid degradation in *C. testosteroni*: degradation of A,B,C,D-rings and removal of the hydroxyl group at C12

The function of ORF7 (*steC*) and ORF6 (*steD*) was revealed in this study and, therefore, the role of all genes in the two steroid degradation clusters on both end of the mega-cluster responsible for steroidal A,B,C,D-ring breakdown was identified. Hereafter, we summarized the entire steroid degradation pathway in *C. testosteroni* TA441 (Fig. 6).

**Fig. 6.**
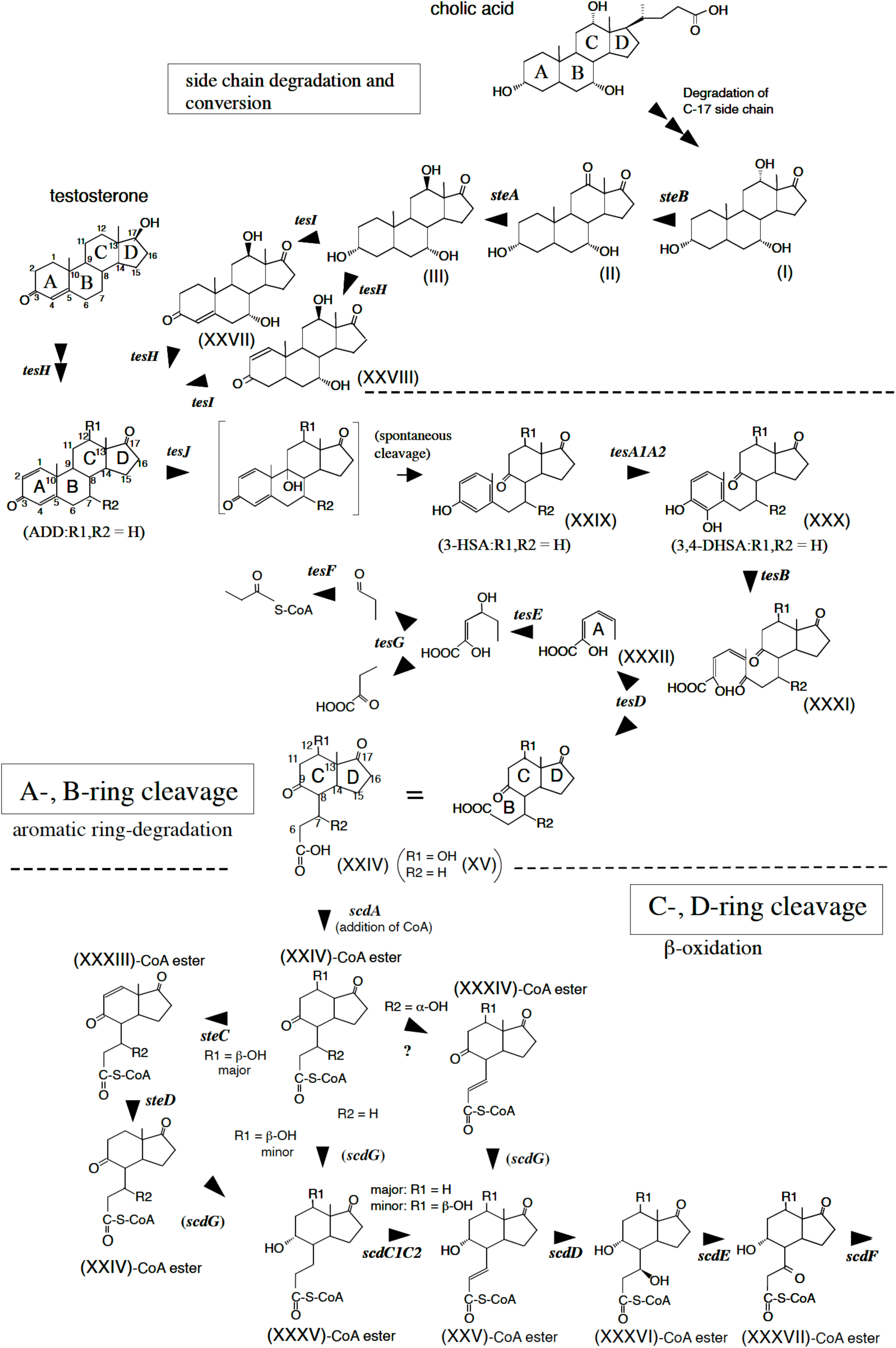
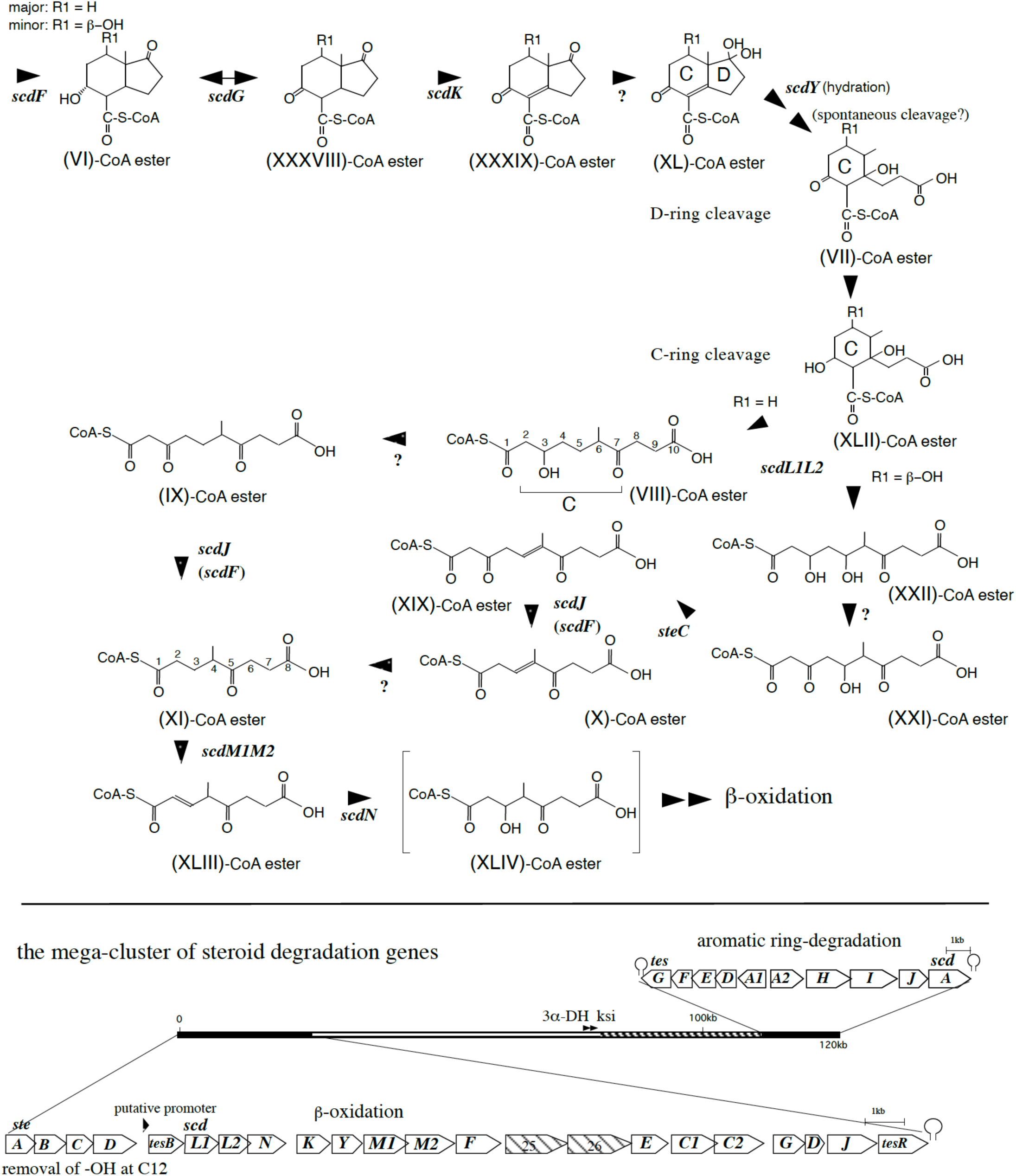
Review of steroid degradation pathway in *Comamonas testosterone* TA441. The mega-cluster of steroid degradation genes in TA441 is shown below the degradation pathway; the aromatic ring-degradation gene cluster (*tesG* to *scdA*) and the β-oxidation gene cluster (*steA* to *tesR*) locate both ends of this 120kb-mega cluster. 3α-Hydroxy-dehydrogenase (3α-DH) gene and 3-ketosteroid Δ4-5 isomerase (ksi) gene are in the DNA region between the two clusters. Possible degradation genes for the side chain at C17 of cholic acid are in the DNA region indicated with stripes. Compounds are; 3α,7α,12α-trihydroxy-17-oxo-androstan, (**I**); 3α,7α-dihydroxy-12,17-oxo-androstan, (**II**); 3α,7α,12β-trihydroxy-17-oxo-androstan, (**III**); 7α,12β-dihydroxy-androst-4-ene-3,17-dione, (**XXVII**); 7α,12β-dihydroxy-androst-1-ene-3,17-dione, (**XXVIII**); androsta-1,4-diene-3,17-dione, (ADD) (R1, R2=H); 3-hydroxy-9,10-secoandrosta-1,3,5(10)-triene-9,17-dione (3-HSA) (R1, R2=H), (**XXIX**); 3,4-dihydroxy-9,10-secoandrosta-1,3,5(10)-triene-9,17-dione (3,4-DHSA) (R1, R2=H), (**XXX**); 4,5-9,10-diseco-3-hydroxy-5,9,17-trioxoandrosta-1(10),2-dien-4-oic acid (R1, R2=H), (**XXXI**); (2*Z*,4*Z*)-2-hydroxyhexa-2,4-dienoic acid, (**XXXII**); 9,17-dioxo-1,2,3,4,10,19-hexanorandrostan-5-oic acid (3aα-*H*-4α [3′-propionic acid]-7aβ-methylhexahydro-1,5-indanedione) (R1, R2=H), (**XXIV**); 12β-hydroxy-9,17-dioxo-1,2,3,4,10,19-hexanorandrostan-5-oic acid (R1=βOH, R2=H), (**XV**); 9,17-dioxo-1,2,3,4,10,19-hexanorandrost-10(12)-en-5-oic acid (R2=H), (**XXXIII**); 9,17-dioxo-1,2,3,4,10,19-hexanorandrost-6-en-5-oic acid (R1=H), (**XXXIV**); 9α-hydroxy-17-oxo-1,2,3,4,10,19-hexanorandrostan-5-oic acid (R1=H), (**XXXV**); 9α-hydroxy-17-oxo-1,2,3,4,10,19-hexanorandrost-6-en-5-oic acid (R1=H), (**XXV**); 7β,9α-dihydroxy-17-oxo-1,2,3,4,10,19-hexanorandrostan-5-oic acid (R1=H), (**XXXVI**); 9α-hydroxy-7,17-dioxo-1,2,3,4,10,19-hexanorandrostan-5-oic acid (R1=H), (**XXXVII**); 9α-hydroxy-17-oxo-1,2,3,4,5,6,10,19-octanorandrostan-7-oic acid (R1=H), (**VI**); 9,17-dioxo-1,2,3,4,5,6,10,19-octanorandrostan-7-oic acid (R1=H), (**XXXVIII**); 9,17-dioxo-1,2,3,4,5,6,10,19-octanorandrost-8(14)-en-7-oic acid (R1=H), (**XXXIX**); 17-dihydroxy-9-oxo-1,2,3,4,5,6,10,19-octanorandrost-8(14)-en-7-oic acid (R1=H), (**XL**); 14-hydroxy-9-oxo-1,2,3,4,5,6,10,19-octanor-13,17-secoandrostane-7,17-dioic acid (R1=H), (**VII**); 9,14-dihydroxy-1,2,3,4,5,6,10,19-octanor-13,17-secoandrostane-7,17-dioic acid (R1=H), (**XLII**); 3-hydroxy-6-methyl-7-oxo-decane-1,10-dioic acid, (**VIII**); methyl-3,7-dioxo-decane-1,10-dioic acid, (**IX**); 3,5-dihydroxy-6-methyl-7-oxo-decane-1,10-dioic acid, (**XXII**); 5-hydroxy-6-methyl-3,7-dioxo-decane-1,10-dioic acid, (**XXI**); 6-methyl-3,7-dioxo-dec-5-ene-1,10-dioic acid, (**XIX**); 4-methyl-5-oxo-oct-3-ene-1,8-dioic acid, (**X**); 4-methyl-5-oxo-octane-1,8-dioic acid, (**XI**); 4-methyl-5-oxo-oct-2-ene-1,8-dioic acid, (**XLIII**); and 3-hydroxy-4-methyl-5-oxo-octane-1,8-dioic acid, (**XLIV**). Enzymes are; SteA (dehydrogenase for 12α-OH to 12-ketone), SteB (hydrogenase for 12-ketone to 12β-OH), TesH (Δ1-dehydrogenase), TesI (Δ4-dehydrogenase), TesJ (ADD**-**hydroxylase at C9), TesA1A2 (3-HSA {**XXIX}-**hydroxylase at C4), TesB (*meta*-cleavage enzyme for 3.4-DHSA {**XXX}**), TesD (**XXXI-**hydrolase), TesE (**XXXII-**hydratase), TesF (aldolase), TesG (acetoaldehyde dehydrogenase), SteC (dehydradase for 12β-OH to produce a double at C10(12)), SteD (reductase for a double at C10(12) to a single bond), ScdA (CoA-transferase for **XXIV**), ScdG (hydrogenase primarily for 9-OH of **VI-**CoA ester), ScdC1C2 (Δ6-dehydrogenase for **XXXV-**CoA ester), ScdD (**XXV-**CoA ester hydratase), ScdE (**XXXVI-**CoA ester dehydrogenase at C7), ScdF (**XXXVII-**CoA ester thiolase/CoA-transferase), ScdK (Δ8(14)-dehydrogenase for **XXXVIII-**CoA ester), ScdY (**XL-**CoA ester hydratase), ScdL1L2 (putative CoA-transferase/isomerase necessary for C-ring cleavage of **XLII-**CoA ester), ScdM1M2 (**XI-**CoA ester dehydrogenase), and ScdN (**XLIII-**CoA ester hydratase). Genes for the C-, D-, and cleaved B-ring degradation are induced by the compounds with steroidal four rings, but are not induced with indane (**XXIV**) and the derivatives (unpublished data).

#### Side chain degradation and conversion

Cholic acid degradation is initiated by the removal of the side chain at position C17. The major intermediates identified in a previous study (29) pointed to β-oxidation as the main degradation mechanism. The corresponding genes were thought to localize to the mega-cluster (striped regions in Fig. 6). After the side chain at C17 is removed, the C12α hydroxyl group on the C-ring (**I**) is converted to a ketone (**II**) by SteA, followed by hydrogenation of the ketone to the C12β hydroxyl group (**III**) by SteB (27). This inversion of stereochemistry is indispensable for subsequent B-ring cleavage.

#### A,B-ring cleavage

After dehydrogenation of the 3β hydroxyl group to a ketone and dehydrogenation of the A-ring by TesH (Δ1 dehydrogenase) and TesI (Δ4 dehydrogenase) to produce androsta-1,4-diene-3,17-dione or one of its derivatives, addition of a hydroxyl group at position C9 by TesJ (formerly ORF17-encoded protein) leads to spontaneous cleavage of the B-ring. This is accompanied by aromatization of the A-ring to 3-hydroxy-9,10-secoandrosta-1,3,5(10)-triene-9,17-dione (R1, R2 = H) (**XXIX**) (30, 31). A hydroxyl group is added at position C4 by TesA1A2 (**XXX**) (32) and the aromatized A-ring is cleaved by the *meta*-cleavage enzyme TesB (**XXXI**) (33), which is followed by TesD-mediated hydrolysis of **XXXI** into cleaved A-ring, (2Z,4Z)-2-hydroxyhexa-2,4-dienoic acid (**XXXII**), and C,D-ring with cleaved B-ring, **XXIV** (34, 35). This process is similar to the bacterial “*meta*-cleavage pathway” responsible for aromatic compound degradation such as biphenyl.

#### C,D-ring cleavage

In contrast to A,B-ring cleavage, C,D-ring cleavage proceeds through a series of β-oxidation cycles. After hydrolysis, CoA is added to **XXIV** at C5 by the CoA transferase ScdA (29). Removal of the C12β hydroxyl group and hydrogenation of the C9 ketone to a C9 hydroxyl occurs prior to the first β-oxidation cycle, which removes the cleaved B-ring. This observation is based on accumulation of 7,12β-dihydroxy-9,17-oxo-1,2,3,4,10,19-hexanorandrostan-5-oic acid (R1 = βOH, R2 = OH) (**XXIV**) in ScdA^-^ incubated with cholic acid; whereas intermediates in the β-oxidation of the cleaved B-ring have a hydroxyl group at C9. Most intermediates in the first β-oxidation cycle do not possess a C12β hydroxyl group, but C,D-ring cleavage can proceed also in its presence, albeit less efficiently. Removal of the C12β hydroxyl group is a two-step reaction that involves dehydration by ScdC and hydrogenation by ScdD. Hydrogenation of the C9 ketone to C9 hydroxyl group is indispensable to initiate β-oxidation, but the corresponding enzyme has not been identified. ScdG can potentially catalyze this reaction, yet the primary function of ScdG is conversion of the C9 hydroxyl group to C9 ketone on 9α-hydroxy-17-oxo-1,2,3,4,5,6,10,19-octanorandrostan-7-oic acid (R1 = H) (**VI**)-CoA ester, a compound generated by the first β-oxidation (36). There may be one or more enzymes other than ScdG, which act on the C9 ketone/hydroxyl group. **XXIV**-CoA ester is converted to 9α-hydroxy-17-oxo-1,2,3,4,10,19-hexanorandrostan-5-oic acid (R1 = H) (**XXXV**)-CoA ester, and then dehydrogenated by ScdC1C2 to **XXV**-CoA ester (37). When a hydroxyl group at C7 is present, **XXV**-CoA ester is produced in ScdC1C2^-^ cultures, suggesting a bypass route (presumably via dehydration) to produce **XXV**-CoA ester (Fig. 6: **XXIV**-CoA ester (R2 = OH) → **XXXIV**-CoA ester → **XXV-**CoA ester) (38). **XXV**-CoA ester undergoes β-oxidation to **VI**-CoA ester via ScdD hydratase, ScdE dehydrogenase, and ScdF CoA-transferase (39). Throughout the β-oxidation of the cleaved B-ring to the cleavage of the C-ring, compounds with “C9 ketone and a double bond at C8(14)” and “C9 hydroxyl group and a single bond at C8(14)” are major in most of the mutant cultures. Given that compounds with “C9 ketone and a single bond at C8(14)” and “C9 hydroxyl group and a double bond at C8(14)” have not been isolated except for **XXIV** and the derivatives, their CoA-esters are likely unstable and are converted to either one of the former two compounds. **VI**-CoA ester is dehydrogenated to 9,17-dioxo-1,2,3,4,5,6,10,19-octanorandrostan-7-oic acid (R1 = H) (**XXXVIII**)-CoA ester; however, this is a reversible reaction and **XXXVIII**-CoA ester is readily dehydrogenated by ScdK at C8(14) to yield 9,17-dioxo-1,2,3,4,5,6,10,19-octanorandrost-8(14)-en-7-oic acid (R1 = H) (**XXXIX**)-CoA ester (36). Next, a water molecule is added at position C17 to produce a geminal diol and the D-ring is cleaved at position C13(17) following addition of another water molecule at C14 by ScdY (accompanying paper). The geminal diol, 17-dihydroxy-9-oxo-1,2,3,4,5,6,10,19-octanorandrost-8(14)-en-7-oic acid (R1 = H) (**XL**), was detected in the present study (accompanying paper), but the enzyme for the production is unclear. CoA-esters of 9-oxo-1,2,3,4,5,6,10,19-octanor-13,17-secoandrost-8(14)-ene-7,17-dioic acid (**XLI**), 9-hydroxy-1,2,3,4,5,6,10,19-octanor-13,17-secoandrosta-13-en-17-oic acid (**XLV**), and 13-hydroxy-9-oxo-1,2,3,4,5,6,10,19-octanor-13,17-secoandrost-8(14)-en-17-oic acid (**XLVI**) were at first proposed as intermediates generated along with the C-ring and cleaved D-ring based on analysis of ScdL1L2^-^ cultures (40). However, subsequent studies suggested **XLI** was produced from 14-hydroxy-9-oxo-1,2,3,4,5,6,10,19-octanor-13,17-secoandrostane-7,17-dioic acid (**VII**)-CoA ester during the isolation procedure (41) and the major product of ScdL1L2 was in fact 3-hydroxy-6-methyl-7-oxo-decane-1,10-dioic acid (R1 = H) (**VIII**)-CoA ester (accompanying paper). **VIII** and **XLV** suggested that the substrate of ScdL1L2 was 9,14-dihydroxy-1,2,3,4,5,6,10,19-octanor-13,17-secoandrostane-7,17-dioic acid (**XLII**)-CoA ester and the function of ScdL1L2 was conversion of **XLII-**CoA ester to **VIII**-CoA ester. Given the similarity with steroid degradation by *M. tuberculosis*, whereby the C-ring of **XLI**-CoA ester is cleaved by the IpdAB hydrolase (a homologue of ScdL1L2) in the presence of FadA6 (a homologue of ScdF) to produce 6-methyl-3,7-dioxo-decane-1,10-dioic acid (R1 = H) (**IX**)-CoA ester (42), the details of the process in *C. testosteroni* and other genera of bacteria may require further elucidation. The ring cleavage product, **VIII**-CoA ester, undergoes the second β-oxidation, whereby the C3 hydroxyl group is dehydrogenated to **IX**-CoA ester, followed by removal of two carbons by the ScdJ CoA-transferase (41). ScdF, and maybe other CoA-transferases, may contribute to this reaction because disruption of ScdJ did not completely stop the conversion of **IX**-CoA ester to 4-methyl-5-oxo-octane-1,8-dioic acid (**XI**)-CoA ester. **XI**-CoA ester undergoes the third β-oxidation involving the ScdM1M2 dehydrogenase, followed by the ScdN hydratase (43), which catalyzes the last reaction among those catalyzed by the enzymes encoded in these two clusters. When the initial steroid compound harbors a C12 hydroxyl group, a portion of it undergoes C,D-ring cleavage with the hydroxyl group, thereby yielding **XXII**-CoA ester, **VIII** derivative with a hydroxyl group at C5 (C12 on steroidal C-ring), and **XI**-CoA ester obtained via 4-methyl-5-oxo-oct-3-ene-1,8-dioic acid (**X**)-CoA ester (43). The low substrate specificity of degradation enzymes allows bacteria to utilize a variety of steroid compounds.

## MATERIALS AND METHODS

### Abbreviations

HPLC, high-performance liquid chromatography; LC/MS, high-performance liquid chromatography-mass spectrometry; RT, retention time; MS, mass spectrometry; HRMS; high-resolution mass spectrometry, MW, molecular weight; CoA, coenzyme A.

### Culture conditions

Mutant strains of *C. testosteroni* TA441 were grown at 30°C in a mixture of equal volumes of Luria-Bertani (LB) medium and C medium (a mineral medium for TA441) (33) with suitable carbon sources. This mixed media is used because the mutants accumulate more amount of intermediate compounds than with C medium or LB medium (unpublished data). Cholic acid and other steroids were added as filter-sterilized DMSO solutions with a final concentration of 0.1% (w/v).

### Construction of gene-disrupted mutants, plasmids, and mutants for complementation experiments

For construction of ORF7^-^6^-^ mutant, (Table 2), pUC19(44)-based plasmid carrying DNA region of TA441 with insertion of a kanamycin-resistance gene (Km^r^) without a terminator between the *Apa*I site in ORF7 and *Bgl*II site in ORF6 (pUCORF76-Km^r^) was used (Table 3). The plasmid was introduced into TA441 via electroporation. The mutants with the insertion by homologous recombination was selected on LB plates with kanamycin (45). Insertion of the Km^r^ was confirmed by southern hybridization and/or PCR-amplification. DNA fragments containing ORF7, ORF6, and ORF76 were obtained by PCR amplification and introduced into a broad-host-range plasmid pMFY42 (28), which can be maintained in *Pseudomonas spp.* and several related species conferring tetracycline resistance to construct pMFYORF7 pMFYORF6, and pMFYORF76, respectively. Retention of the plasmids by the gene-disrupted mutants and the transformants was confirmed by PCR-amplification. For PCR amplification, DNA polymerase KOD -plus- ver. 2 (TOYOBO, Japan) was used. The primers are listed in Table 4.

### Isolation and identification of compounds accumulated in ORF6^-^ culture

ORF6^-^ was grown in 500 ml of 1/2LB+C medium with 0.1% cholic acid and incubated at 30 °C for about 3 days. 6b and 6c were isolated the amount of 10mg and 4.1mg, respectively, from this culture. The amount of 6a was too small, so ORF6^-^ was grown again in total of 1000 ml of 1/2LB+C medium for 2 days and 6mg of 6a was isolated. After the incubation, the culture was extracted twice with the same volume of ethyl acetate. The ethyl acetate fraction was dried and concentrated, dissolved in a small amount of methanol. The compounds were separated by Waters 600 HPLC (Nihon Waters, Tokyo, Japan) with an Inertsil ODS-3 column (20 x 250 mm, GL Science) and a solvent, with the composition CH_3_CN:CH_3_OH:H_2_O:TFA of 50:10:40:0.05, flow rate 1 ml/min, at 40 °C. and the fraction containing each compound was collected from the eluent. The fraction was dried and kept in the refrigerator and subjected to FAB-MS and NMR analyses.

### General experimental procedures

FAB-MS (positive-ion mode) was recorded on a JEOL JMS-700 mass spectrometer (JEOL Ltd., Tokyo, Japan), using a glycerin matrix. 1D and 2D NMR spectra were recorded on a JNM-ECP500 or JNM-ECA600 spectrometer (JEOL Ltd, Tokyo, Japan). Tetramethylsilane (TMS) at 0 ppm in CDCl_3_ solution and residual proton signal at 2.49 ppm in DMSO-d6 solution were used as internal references for ^1^H chemical shifts. ^13^C chemical shifts were obtained with reference to DMSO-d6 (39.5 ppm) or CDCl_3_ (77.0 ppm) at 25°C.

### HPLC analysis (Fig. 2)

After addition of a double volume of methanol to the culture, the mixture was centrifuged, and the supernatant was directly injected into an HPLC. The HPLC (Alliance 2695 with UV detector and 996 photodiode array detector, Nihon Waters, Tokyo, Japan) equipped with an Inertsil ODS-3 column (4.6 × 250 mm, GL Sciences Inc., Tokyo, Japan) was used, and elution was carried out using a linear gradient from 20% solution A (CH_3_CN:CH_3_OH:TFA = 95:5:0.05) and 80% solution B (H_2_O:CH_3_OH:TFA = 95:5:0.05) to 65% solution A and 35% solution B over 10 mins; this was maintained for 3 mins, and then changed to 20% solution A. The flow rate was 1.0 mL/min.

### Reverse-phase liquid chromatography with tandem mass spectrometry (LC/MS/MS)

For LC/MS/MS analysis, 500μl culture was acidified with 6N HCl and extracted with ethyl acetate. After ethyl acetate was vaporized at room temperature, 600μl methanol was added, stored at -80°C and subjected to analysis within a day. 2 µl of the samples was injected into the Triple Quadrupole Mass Spectrometry. Agilent 1100 HPLC (Agilent, CA) with a mass spectrometer (Applied Biosystems 4000 Q-TRAP, MS) was used with L-column2 ODS (1.5 × 150 mm) Type L2-C 18.5µm, 12mm (GL Science, Tokyo, Japan) and elution was carried out using 90% solution C (H_2_O:HCOOH = 100:0.1) and 10% solution A for 1min, followed by a linear gradient from 90% solution C and 10% solution A to 20% solution C and 80% solution A over 7 min, which was maintained for 2 min. The flow rate was 0.2 ml/min. The desolvation temperature for the mass spectrometer was fixed at 450 °C. The collision energy was 20 V.

### Data availability

The authors agree that any materials and data that are reasonably requested by others are available from a publicly accessible collection or will be made available in a timely fashion, at reasonable cost, and in limited quantities to members of the scientific community for noncommercial purposes.

## Acknowledgments

MH appreciates Dr. Reizo Kato (RIKEN) and Dr. Yousoo Kim (head of Surface and Interface Science Laboratory, RIKEN) for thoughtful supports and advice. MH thanks Dr. Takemichi Nakamura (Molecular Structure Characterization Unit, RIKEN CSRS, WAKO) for his assistance in mass spectrometry and Dr. Hiroyuki Koshino (Molecular Structure Characterization Unit, RIKEN CSRS, WAKO) for his assistance in NMR analysis.

## FIGURE LEGENDS (SUPPLEMENTARY MATERIAL)

Fig S1 LC/MS analysis of the culture of the ORF6^-^ mutant incubated with cholic acid (S1-1), the ORF7^-^ mutant with cholic acid(S1-2), deoxycholic acid (S1-3), and chenodeoxycholic acid (S1-4). Closed arrows indicate the peak of the compound presented on the left side of the columns. Compounds are; 12β-hydroxy-9,17-dioxo-1,2,3,4,10,19-hexanorandrostan-5-oic acid, (**XV**); 12β,17-dihydroxy-9-oxo-1,2,3,4,10,19-hexanorandrostan-5-oic acid, (**XVI**); 7β,9α-dihydroxy-17-oxo-1,2,3,4,10,19-hexanorandrostan-5-oic acid, (**XXXVI**); 7α,12β-dihydroxy-9,17-dioxo-1,2,3,4,10,19-hexanorandrostan-5-oic acid, (**XXIVa**); 9,17-dioxo-1,2,3,4,10,19-hexanorandrostan-5-oic acid, (**XXIV**); 9α-hydroxy-17-oxo-1,2,3,4,10,19-hexanorandrost-6-en-5-oic acid, (**XXV**); 17-hydroxy-9-oxo-1,2,3,4,10,19-hexanorandrost-10(12)-en-5-oic acid, (**XXVI**); 9α-hydroxy-17-oxo-1,2,3,4,10,19-hexanorandrosta-6,10(12)-dien-5-oic acid, (**XVII**); 9α-hydroxy-17-oxo-1,2,3,4,5,6,10,19-octanorandrostan-7-oic acid, (**VI**); 9α,12β-dihydroxy-17-oxo-1,2,3,4,5,6,10,19-octanorandrostan-7-oic acid, (**VIa**); 9-oxo-1,2,3,4,5,6,10,19-octanor-13,17-secoandrost-8(14)-ene-7,17-dioic acid, (**XLI**); 6-methyl-3,7-dioxo-dec-5-ene-1,10-dioic acid, (**XIX**); 5-hydroxy-6-methyl-3,7-dioxo-decane-1,10-dioic acid, (**XXI**); 3-hydroxy-6-methyl-7-oxo-dec-5-ene-1,10-dioic acid, (**XX**); 3,5-dihydroxy-6-methyl-7-oxo-decane-1,10-dioic acid, (**XXII**); 4-methyl-5-oxo-octane-1,8-dioic acid, (**XI**); 9,17-dioxo-1,2,3,4,10,19-hexanorandrost-10(12)-en-5-oic acid, (**XXXIII**); and 9,17-dioxo-1,2,3,4,10,19-hexanorandrost-6-en-5-oic acid, (**XIII**). Mass spectrums of these compounds are presented in Fig. S2 in Supplementary Material. Vertical axis indicates intensity (count/sec) and the horizontal axis indicates RT (min).

Fig S2 The mass spectrums of compounds detected in the culture of the ORF6^-^ mutant and the ORF7^-^ mutant analyzed by LC/MS/MS. Alphabets on the columns correspond to those in Fig. S1. Compounds do not appear in Fig. S1 but have the same parental peak (*m/z*) are given the same alphabet. Compounds with * were identified in the previous study. Compounds are; A1, 12β-hydroxy-9,17-dioxo-1,2,3,4,10,19-hexanorandrostan-5-oic acid (**XV**); A2, 9α-hydroxy-7,17-dioxo-1,2,3,4,10,19-hexanorandrostan-5-oic acid, (**XXXVII**)*; B1, 12β,17-dihydroxy-9-oxo-1,2,3,4,10,19-hexanorandrostan-5-oic acid, (**XVI**); B2, 7β,9α-dihydroxy-17-oxo-1,2,3,4,10,19-hexanorandrostan-5-oic acid, (**XXXVI**)*; C, 7α,12β-dihydroxy-9,17-dioxo-1,2,3,4,10,19-hexanorandrostan-5-oic acid, (**XXIVa**); D1, 17-hydroxy-9-oxo-1,2,3,4,10,19-hexanorandrost-10(12)-en-5-oic acid, (**XXVI**); D2, 9α-hydroxy-17-oxo-1,2,3,4,10,19-hexanorandrost-6-en-5-oic acid (R1=H), (**XXV**)*; D3, 9,17-dioxo-1,2,3,4,10,19-hexanorandrostan-5-oic acid, (**XXIV**)*; E1, fragment of **XV**; E2, 17-hydroxy-9-oxo-1,2,3,4,10,19-hexanorandrost-6,10-dien-5-oic acid, (**XII**); E3&5, 9α-hydroxy-17-oxo-1,2,3,4,10,19-hexanorandrosta-6,10(12)-dien-5-oic acid (**XVII**) stereo isomers?; E4, 9,17-dioxo-1,2,3,4,10,19-hexanorandrost-6-en-5-oic acid, (**XIII**); E6, 9,17-dioxo-1,2,3,4,10,19-hexanorandrost-11-en-5-oic acid (**XXIII**); E7, lacton form of 12-hydroxy-9,17-dioxo-1,2,3,4,10,19-hexanorandrost-6-en-5-oic acid; F, 9α-hydroxy-17-oxo-1,2,3,4,5,6,10,19-octanorandrostan-7-oic acid (**VI**)*; G1, 9α,12β-dihydroxy-17-oxo-1,2,3,4,5,6,10,19-octanorandrostan-7-oic acid, (**VIa**); G2, 9α-hydroxy-1,2,3,4,10,19-hexanorandrost-10-en-5-oic acid ?; G3, 9α-hydroxy-1,2,3,4,10,19-hexanorandrost-13-en-5-oic acid, (**VLV**)*; G4&G5, intermediate compounds produced before addition of CoA at C5; H1, decarboxylated derivative of 9-oxo-1,2,3,4,5,6,10,19-octanor-13,17-secoandrost-8(14)-ene-7,17-dioic acid (**XLI**)*, H2, a derivative of 9α,12β-dihydroxy-1,2,3,4,10,19-hexanorandrost-13-en-5-oic acid; I1, decarboxylated derivative of 13-hydroxy-9-oxo-1,2,3,4,5,6,10,19-octanor-13,17-secoandrost-8(14)-en-17-oic acid (**XLVI**)*; I2, decarboxylated derivative of 12-hydroxy-9-oxo-1,2,3,4,5,6,10,19-octanor-13,17-secoandrost-8(14)-en-17-oic acid; J1, 13-hydroxy-9-oxo-1,2,3,4,5,6,10,19-octanor-13,17-secoandrost-8(14)-en-7,17-dioic acid (**XLVI**); J2&J3, stereo isomers of 12-hydroxy-9-oxo-1,2,3,4,5,6,10,19-octanor-13,17-secoandrost-8(14)-en-7,17-dioic acid ?; J4, 6-methyl-3,7-dioxo-dec-5-ene-1,10-dioic acid (**XIX**); K1, 5-hydroxy-6-methyl-3,7-dioxo-decane-1,10-dioic acid (**XXI**); K2, an intermediate compound produced before addition of CoA at C5; L, methyl-3,7-dioxo-decane-1,10-dioic acid-CoA ester,(**VIII**); M, 3-hydroxy-6-methyl-7-oxo-dec-5-ene-1,10-dioic acid (**XX**); and N, 4-methyl-5-oxo-octane-1,8-dioic acid (**XI**)*. The vertical axis indicates relative intensity (%) and the horizontal axis indicates mass (*m/z*) in mass spectra. In mass chromatogram, the vertical axis indicates intensity (count/sec) and the horizontal axis indicates RT (min).

Fig S3 Additional data for Fig. 4. Complementation experiments with ORF7^-^, ORF6^-^, and ORF7^-^6^-^ mutants. The mass chromatograms of each mutant culture incubated with 0.1% cholic acid for 7days are shown. Mutants are; ORF7^-^ mutant carrying pMFY42 (ORF7^-^ with pMFY42) (negative control) (A), carrying pMFYORF7 (pMFY42 carrying ORF7) (B), ORF6^-^ mutant carrying pMFY42 (ORF6^-^ with pMFY42) (negative control) (C), carrying pMFYORF6 (ORF6^-^ with pMFYORF6) (D), ORF7^-^6^-^ mutant carrying pMFY42 (ORF7^-^6^-^ with pMFY42) (negative control) (F), carrying pMFYORF7 (G), and carrying pMFYORF76 (H). Mass chromatograms of *m/z* 227 (S3-1), *m/z* 241 (S3-2), *m/z* 197 (S3-3), and *m/z* 201 (S3-4) are shown. Compounds are; 9α,12β-dihydroxy-17-oxo-1,2,3,4,5,6,10,19-octanorandrostan-7-oic acid (**VIa**), 6-methyl-3,7-dioxo-dec-5-ene-1,10-dioic acid (**XIX**), 5-hydroxy-6-methyl-3,7-dioxo-decane-1,10-dioic acid (**XXI**), 13-hydroxy-9-oxo-1,2,3,4,5,6,10,19-octanor-13,17-secoandrost-8(14)-en-17-oic acid (**VLVI**), and 4-methyl-5-oxo-octane-1,8-dioic acid (**XI**). The vertical axis indicates intensity (count/sec) and the horizontal axis indicates RT (min).

